# A modular multi-color fluorescence microscope for simultaneous tracking of cellular activity and behavior

**DOI:** 10.1101/2025.04.01.646316

**Authors:** Euphrasie Ramahefarivo, Leonard Böger, Takkasila Saichol, Behzad Shomali, Luis Alvarez, Monika Scholz

## Abstract

We present a modular epifluorescence tracking microscope which enables ratiometric imaging of muscles, neurons, and other structures in moving animals. The microscope is assembled entirely from commercial parts within 3 hours, making the system broadly accessible. Leveraging the improved brightness and bleaching characteristics of recent genetically-encoded indicators and fluorophores, the simple microscope is even suitable for simultaneous calcium imaging of neurons and behavior as we demonstrate in *C. elegans*. We also show how muscle dynamics in *D. melanogaster* larvae can be analyzed and how dual color fluorescence tracking elucidates inter-species interactions by visualizing both predatory nematodes and their prey. Finally, we showcase a configuration for bright-field imaging, by tracking tardigrade gait as an example of utility for non-model species. The affordability of the hardware and ease of use of the accompanying software make this a suitable tool for education in addition to its use in research.

## Introduction

Access to automated, robust behavioral tracking has transformed research in multiple diverse research areas such as neuroscience, behavioral ecology, and ethology ^1–4^. Combining simple static cameras for high-speed videography with user-friendly deep-learning tools has enabled researchers working to rapidly expand their capabilities to study behaviors and scale up the analysis to many recordings^5–8^. However, smaller animals with sizes ranging from sub-mm (hydra, tardigrades) to multi-mm range such as the nematode *C. elegans*, larval zebrafish, *Drosophila* larvae, or planaria, are typically too small for standard videography and require microscopes for imaging, resulting in small fields of view that do not allow for unrestrained exploration. To this end, tracking microscopes that continually follow an animal as it moves have been employed, keeping it centered in a camera’s field of view (FOV). Tracking microscopy can extend the duration and spatial range over which an animal can move within an experiment beyond setups with static cameras. The resulting videos have the same resolution and the distance from the animal to the camera can remain constant even if animals move far from the starting location, easing downstream analyses.

Brightfield tracking microscopes for animals such as *C. elegans* have a long history, with the first automated worm trackers published already in the early 2000s designed for automated phenotyping of posture and locomotion ^1,3,4,9,10^. Recent setups have improved speed, ease-of-use, and analysis capabilities^4^, detecting additional behaviors like egg-laying and feeding ^11^. Extending these brightfield trackers to detect fluorescent molecules allows extracting additional information, for example, connecting posture to muscle activity ^12^, mapping neuronal activity to behavior ^13–17^, or allowing multi-animal identification ^18^.

In contrast to brightfield tracking microscopes, fluorescence microscopes are often built for specific tasks. Their design requires custom components or expensive hardware, making them inaccessible to a broader group of researchers. The development of novel fluorophores^19–21^ and complementary metal-oxide-semiconductor (CMOS) imaging chips has ushered in a transformative era in fluorescence microscopy, substantially altering the speed of acquisition and the price for cameras suitable for fluorescence measurements^22^. These developments enable fluorescence detection from samples even with weaker labeling and using lower magnification, resulting in larger FOV^23,24^. Additionally, the fast read-out speeds of CMOS cameras enable tracking and recording at high-speed fluorescent cellular structures or organs as the animal moves.

Non-experts often face barriers when accessing newly developed methods due to manufacturing complexity or cost restrictions. To reduce costs, some methods use low-cost 3D-printed parts and custom electronics to replace custom-machined elements or moving elements such as stages ^10,25,26^. However, this approach requires some experience in additive manufacturing and can mean a loss of structural integrity and stability. We therefore set out to develop a dual-color epi-fluorescence tracking microscope purely from commercially available off-the-shelf components.

In this paper, we present a design for a modular epifluorescence tracking microscope suitable for mm-sized animals. Using calcium-imaging in muscles and neurons, we demonstrate its capability for behavioral neuroscience. By imaging predatory interactions between two species of nematodes, we illustrate its use for behavioral ecology. In addition, we demonstrate how the same components can be assembled into a simpler single-color variant of the microscope, and we illustrate the capabilities of this setup for posture detection in a non-model species: the tardigrade. The microscope’s affordability, versatility, and user-friendly design make it a valuable addition to the toolkit of researchers studying small millimeter-scale organisms, offering novel capabilities for fluorescence tracking, all without the need for specialized fabrication.

## Results

### Hardware: Modular design enables multiple imaging modes

We designed a modular epi-fluorescence tracking microscope for small model animal tracking (Fig. 1A-C). The hardware is modular, supporting single-color, dual-color, and brightfield imaging modes, with magnifications from 1 - 4.2x suitable for macro-imaging of whole animals, tissues within living animals, and single neurons (Fig. 1D). The list of required parts is available in Table S2, as well as on the accompanying website. The microscope compensates for animal motion by moving the light source and camera, keeping the sample stationary to reduce vibration and allow for additional components like heaters, coolers, or optogenetic tools. The sample remains in a fixed arena, simplifying the creation of spatially-dependent environments for experiments such as chemotaxis or thermotaxis^4^.

**Figure 1:**
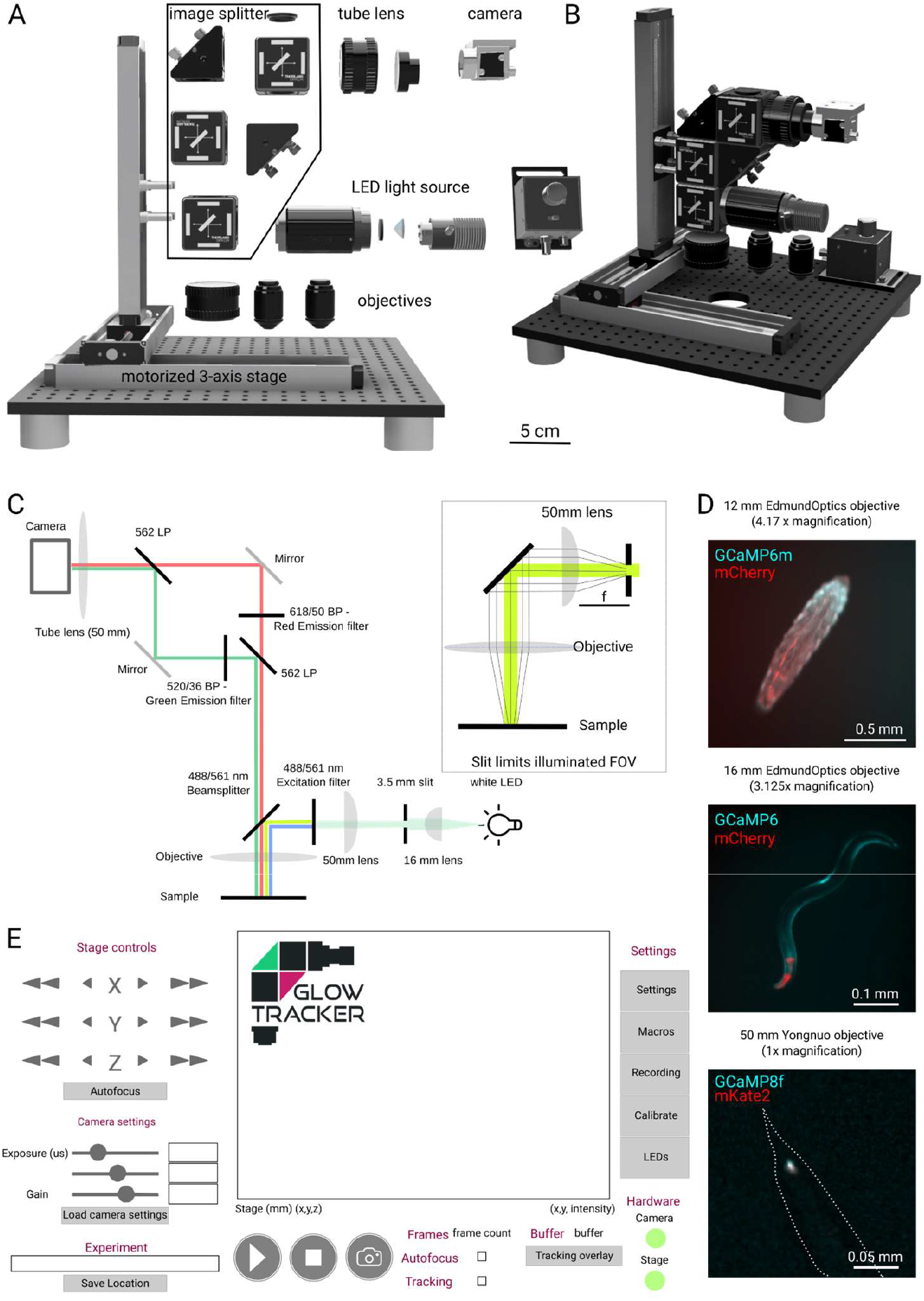
Design of a modular epi-fluorescence tracking microscope. (A) Components and (B) 3D rendering of the dual-color epi-fluorescence tracking microscope (‘macroscope’). (C) Lightpath of the dual color tracking microscope using a slit imaged onto the sample plane to restrict illumination (inset) and a compact image splitter design. (D) Example images of dual color imaging at different magnifications of a Drosophila 3rd instar larvae with muscles expressing mCherry and GCaMP6 at 1x magnification; Image of an adult *C. elegans* with pharyngeal (red) and body wall muscles (cyan) labeled with mCherry and GCaMP6 at 2.78x magnification and *C. elegans* touch receptor neuron PLM labeled with mKate2 (red) and GCaMP8f (cyan) imaged at a magnification of 4.17x. The outline shows the tail of the animal (dashed). Magnification was adjusted by using objectives with a focal length of 50, 18, and 12 mm, respectively. The same 50 mm tube lens was used for all recordings. (E) Schematic of the GlowTracker GUI showing the interface and different features such as calibration and autofocus.

The dual-color configuration projects two-color images side-by-side onto the camera using a compact image splitter in the W-design^27^ (Fig. 1C). For convenience, this feature is matched in the user interface by a corresponding automated color calibration that allows saving the resulting images as registered stacks which are already processed to have the correct overlap between each channel. The two other configurations (brightfield, single color fluorescence) have simplified lightpaths and primarily use a subset of the components of the dual color microscope, allowing users to switch between different use cases (Supplementary Fig. 2, Supplementary Table 1).

**Figure 2:**
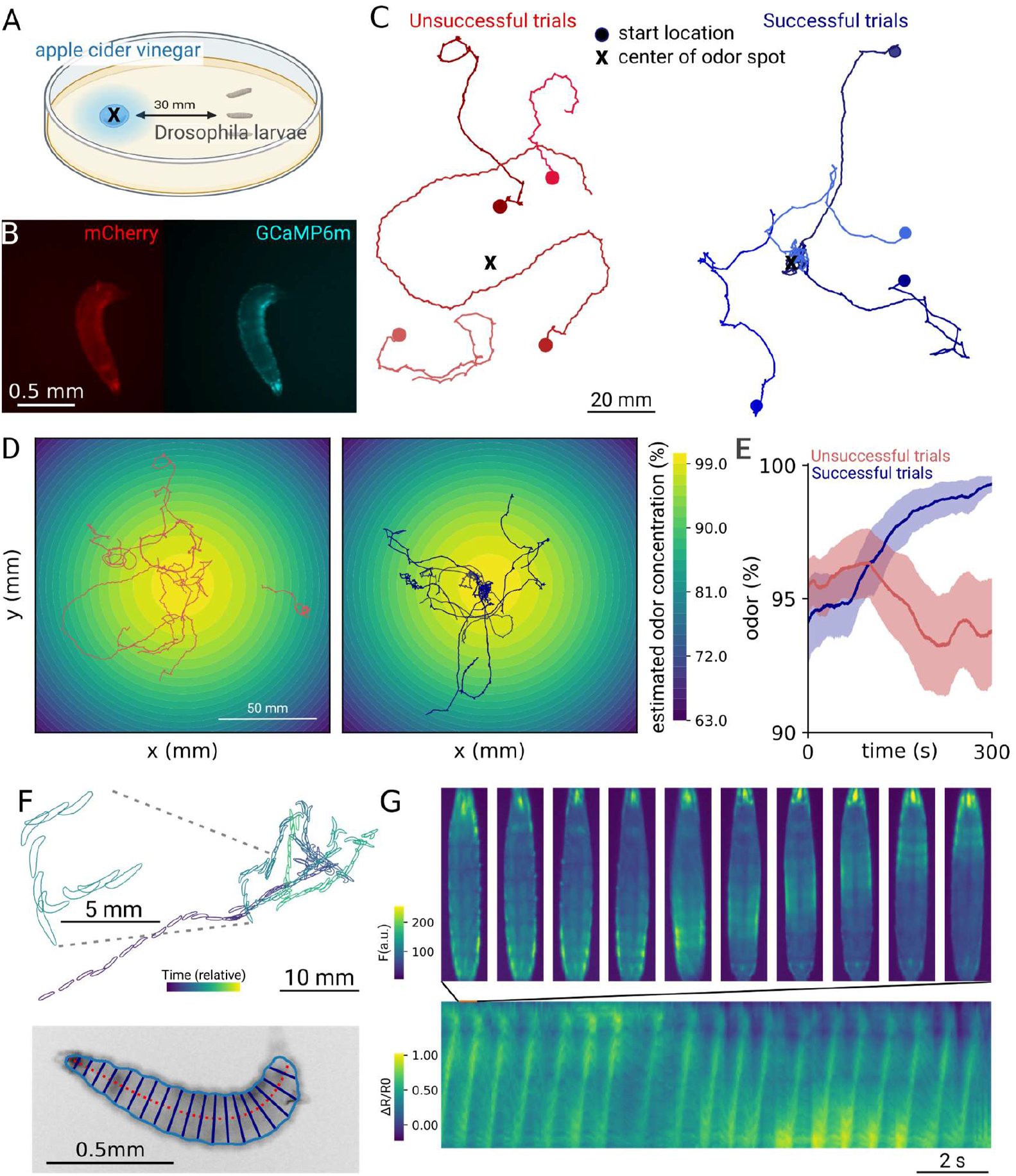
Tracking behavior of *Larvae* while navigating an odor gradient. (A) Schematic of the odortaxis assay. (B) Example images of the dual-view ratiometric calcium imaging during the experiment. (C) Representative example tracks for both successful (blue) and unsuccessful (red) trials. Trials were counted as successful if the larvae entered and remained within 5 mm of the center of the target spot within the 5 minutes of recording time (N=4 selected traces). (D) Estimated odor gradient from a diffusion model of vinegar and (E) odors experienced by the successful (blue) and unsuccessful (red) larvae over time. (F) Example track of a successful larva with the posture overlaid. Posture was extracted from images as shown in the inset below. (G) Kymograph of muscle GCaMP activity for the animal in (F) during straight navigation. The Ca^2+^ waves along the animal body during muscle contraction can be observed. Exemplary images during a single stride are shown on the top.

The magnification of the microscope can be adapted by exchanging the objective (Fig. 1C-E). The standard configuration with the largest FOV results in a 1x magnification and 2.34 µm/px resolution while allowing a 7.4 mm x 5.0 mm FOV. We present three configurations, with resulting magnifications ranging from 1x to 4.2x (Supplementary Table 1). Other dry objectives already present in the lab can be used, requiring only the appropriate adapters (Fig. 1E; see supplementary information to calculate the resulting magnification). The light sources are commercial LEDs which can be matched to the desired fluorophore, such as deep IR for behavioral imaging or a general purpose white LED, which we use throughout due to its broad applicability for many different fluorophores.

The stage is critical for the function of this microscope, as it will determine at which speeds animals can be tracked successfully. To allow easy integration with any computer and operating system, we use a USB 3.0 stage with integrated motor controllers, capable of moving up to 20 mm/s, and a 150 mm travel range, which is sufficient for typical experiments. The camera is a Basler sCMOS model capturing up to 57 full frames per second, featuring a USB 3.0 connection for platform-independent integration without frame grabbers. All hardware components are available from commercial vendors and do not require soldering or manufacturing; only assembly. Detailed documentation on assembly and usage is provided online (see Data and Code availability section), allowing users to assemble the microscope within 3 hours using the illustrated guide (https://scholz-lab.github.io/GlowTracker/).

### Platform-independent software for effective interaction with the hardware

To allow users to interact and perform experiments with the microscope, we have built GlowTracker: an application with a graphical user interface that ties the hardware together. The application can record and display images from the camera, control the stage, and perform automated object tracking (Supplementary Video 1). GlowTracker is built on top of the Python-based Kivy App framework and has been designed to work on Windows, Linux, and macOS (see code availability section for a link to the software). Key capabilities include an autocalibration where the pixel size and stage orientation are automatically determined, and a color calibration feature, which calculates the optimal overlay of multiple color channels to minimize post-processing of the resulting images. In addition, the software includes a macro language, allowing basic scripts to move in a defined area or take z-stacks. Taken together, these features enable flexible usage of the microscope even for users inexperienced in coding.

To determine the GlowTracker GUI’s performance, we ran benchmark tests on image analysis and tracking. We found that tracking performance was most influenced by the exposure time, as this limits the framerate at which images are incoming. In practice, we could obtain image acquisition frame rates of 16 Hz up to 50 Hz for bright samples, which was only limited by the required exposure times (Supplementary Fig.1). To avoid calculating the displacement on blurred frames, we ignore frames where the stage moved in tracking calculations. In addition, the built-in stage controllers have a communication latency of about 25 ms, which is comparable to typical exposure times (Supplementary Fig.1). If faster tracking is desired, users could upgrade to standalone controllers that have much lower communication latencies. However, we found that for a variety of applications the communication speed was not a limiting factor for tracking performance. Depending on the camera frame rate, the system can track (i.e. update the stage position) at 6.5 Hz to 11 Hz. The full description of benchmarking and hardware requirements are discussed in the Supplementary Materials.

In summary, we designed a modular microscope suitable for imaging samples in the few-mm to µm range that allows single- or multi-color imaging. To test the applicability of the setup, we present multiple typical use cases: ratiometric calcium imaging of neurons and muscles of unrestrained animals, tracking of non-labelled model species, as well as visualization of inter-species predatory interactions.

### Simultaneous tracking of posture, muscle activity, and locomotion in *D. melanogaster* larvae

With a speed of about 1 mm/s, Drosophila larvae can explore arenas of multiple tens of cm within minutes, requiring a tracking microscope to bridge the scale between muscle contraction and large-range locomotion ^28^. Fly larvae range between 1.4 mm at their first larval stage up to 3-5 mm at their third larval stage^29^, making them suitable for imaging with the 1x microscope (Fig. 2A,B). Larvae are effective at finding attractive odor sources, using a navigational algorithm that includes active sampling ^30^. As larvae move by peristaltic traveling waves along their anterior-posterior axis interspersed with turns, the navigation is well-described by specific muscle contraction patterns ^31–33^. For example, while traveling straight requires symmetric left-right muscle contractions, turns and bends are created by asymmetric contractions. We aimed to connect the biomechanics of the body to the navigational strategy by simultaneously imaging the larvaes’ navigation along a vinegar gradient and monitoring muscle activity with a genetically-encoded calcium indicator (Fig. 2A-B).

We find that some larvae do not reach the odor spot within the allotted time, despite having sufficient speed to be able to reach the spot (Fig. 2C-E). We confirm that animal size correlates with speed, as had been reported previously ^34^, (Supplementary Fig.4A-C), however, speed did not differ significantly between successful and unsuccessful larvae (Supplementary Fig. 4B). To investigate potential differences in navigational strategies between successful and unsuccessful larvae at the biomechanical level, we analyzed their posture and calculated a virtually straightened animal. Using the two-color imaging, we can ratiometrically correct the GCaMP images, and extract features like fluorescence intensity along the anterior-posterior axis (Fig. 2G), showing characteristic peristaltic waves used for propagation. Similar analyses have enabled the prediction of larval locomotion from muscle activity ^12^, but the amount of motion was constrained by the field of view. Allowing imaging of unrestrained animals in large arenas will be useful for a variety of applications: From basic biomechanics research into how posture translates into movement, to studies of movement disorders. While fictive locomotion assays and assays using non-tracking microscopes have already provided insight ^35,36^, experiments on the microscope can be done in structured environments, such as gradients, and allow for many more strides per larva. In summary, we show that the modular tracking microscope is suitable for tracking larval Drosophila across cm-scaled environments while extracting relevant physiological data.

### Ratiometric imaging of single neurons in moving animals

To demonstrate the dual-color capability in a smaller sample under challenging conditions, we aimed to image single neurons in unrestrained animals. Recent advances in single-neuron labeling, such as the use of Gal4 driver lines ^37^ and newly developed sensory-specific strains ^38^, have allowed for more precise labeling. In addition, fast and bright calcium indicators are now amenable to low-magnification imaging with our microscope. We recently adapted the new generation calcium indicator GCaMP8f for use in nematodes and demonstrated its properties in restrained animals^19^. Using this indicator, we chose to image the dynamics of *C. elegans* touch receptor neurons (TRNs), which induce an escape response when stimulated (Fig. 3A,B). As immobilization has been shown to alter neural activity ^14,35^, we focused on capturing neural responses in a natural, unrestrained environment. Prior work used either custom microscopes ^11,39^, or low-resolution large field-of-view imaging methods in small arenas ^23^. We employed ratiometric imaging techniques to reduce motion artifacts and correct for focusing issues, while enabling high-frame-rate imaging of TRNs. In our experiment, we stimulated *C. elegans* worms with a mechanical vibration for 1s, resulting in the activation of all TRNs ^40^. We tracked the PLM neuron, a TRN located in the tail of *C. elegans*, proposed to control forward escape responses^41,42^. Expectedly, the animals showed increased speed and a significant rise in the PLM neuron’s calcium signal dR/R0 (Fig. 3C-D). Interestingly, when comparing worms that showed reversals within 2 seconds after stimulus onset to those that did not, we could observe distinct calcium traces (Fig. 3E). Worms showing a reversal escape response had a decreased peak activity in the PLM neurons and decayed faster compared to animals who performed a forward escape (Fig. 3F). The microscope is therefore suitable for imaging challenging, small samples in moving animals while allowing for robust tracking and motion correction.

**Figure 3:**
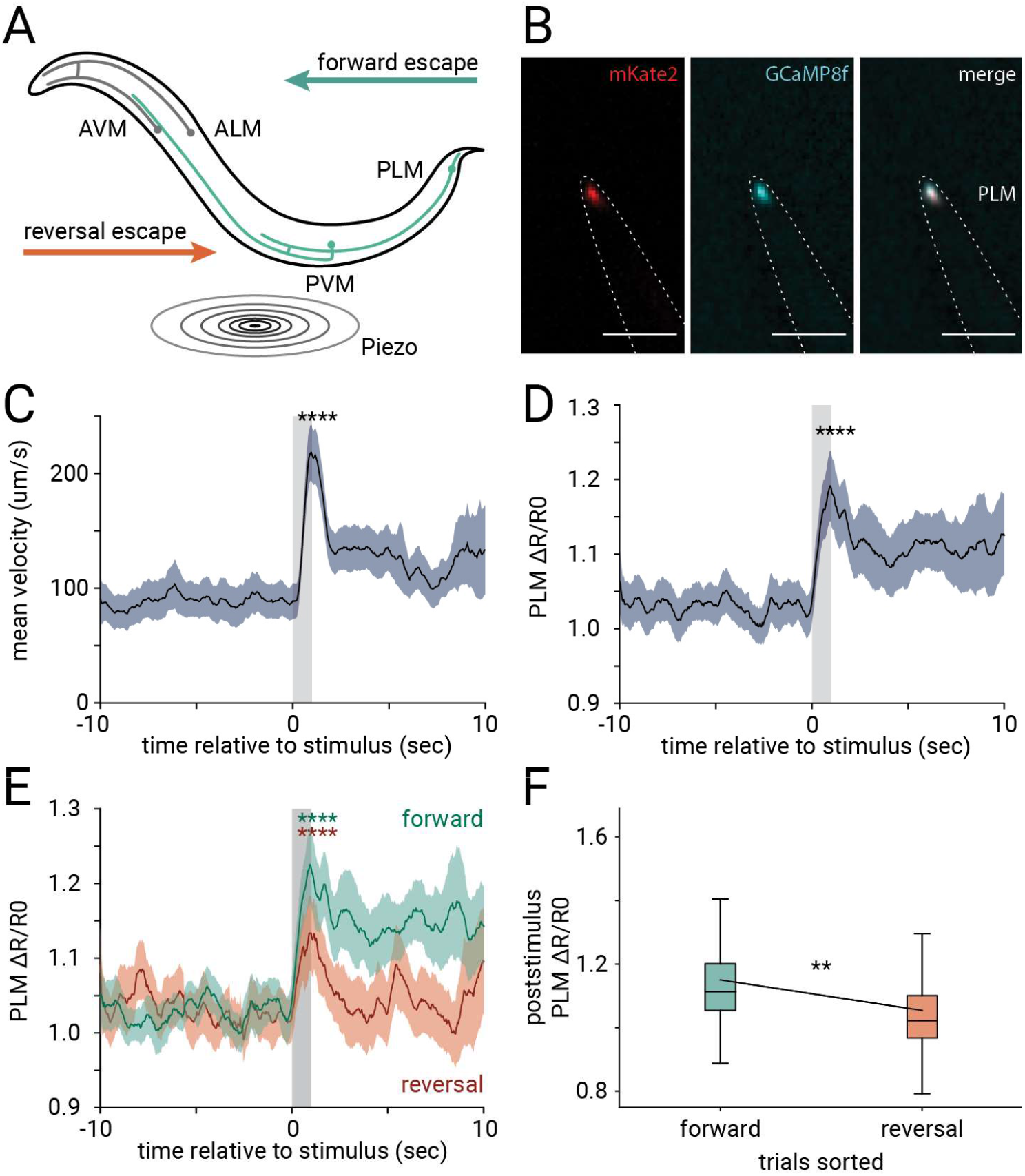
Ratiometric imaging of touch-receptor neurons in unrestrained animals. (A) Schematic of touch receptor neurons in *C. elegans* and behavioral response to gentle mechanical stimulus (B) Touch receptor neuron PLM labeled with GCaMP8f and mKate2 (cyan, red, respectively). Dashed line shows worm outline. Scale bar corresponds to 50 µm. (C) Speed response (mean ± s.e.m.) to light touch stimuli (gray bar) Statistics performed with paired t-test, * p < 0.05 **** p < 0.0001 (D) Neuronal response (mean ± s.e.m.) to light touch stimuli (gray bar). Statistics as in C (E) Neuronal response to light touch stimuli (gray bar) as in E, for trials with reversal (orange) and forward response (green) within 2 sec after stimulus onset. Statistics as in C. (F) Boxplot showing mean poststimulus GCaMP8 fluorescence signal, color as in E. For forward responding worms N=15 worms, n=41 traces, with 1-5 repetitions per animal. The box spans the first to third quartile, with a line marking the median. Error bars extend to the furthest point within 1.5 times the interquartile range. Outliers are excluded. For reversal responding worms N=12 worms, n=20 traces, with 1-3 repetitions per animal. Statistics performed with a mixed linear model, ** p < 0.01.

### Simultaneous imaging of animals and the environment

To study interspecies interactions, we employed the dual-color microscope to visualize predatory behavior between two species of nematodes, the predator *Pristionchus pacificus* and its prey, larval *Caenorhabditis elegans* ^43,44^. *P. pacificus* can prey on non-kin larvae of other nematodes, exhibiting contact-mediated kin detection^45,46^. We tracked *P. pacificus* expressing red fluorescent protein (RFP) in its pharynx ^44^ and detected prey contact using a *C. elegans* strain expressing GFP in its body wall (Fig. 4A). We previously developed a machine-learning model to identify behavioral states of the predator using only the images of its pharynx, however, with the dual-color images we are now able to confirm the model predictions, by also visualizing prey contact directly. The model is able to identify different behavioral states related to feeding, such as predatory biting, as well as non-predatory movement behaviors (roaming, dwelling).

**Figure 4:**
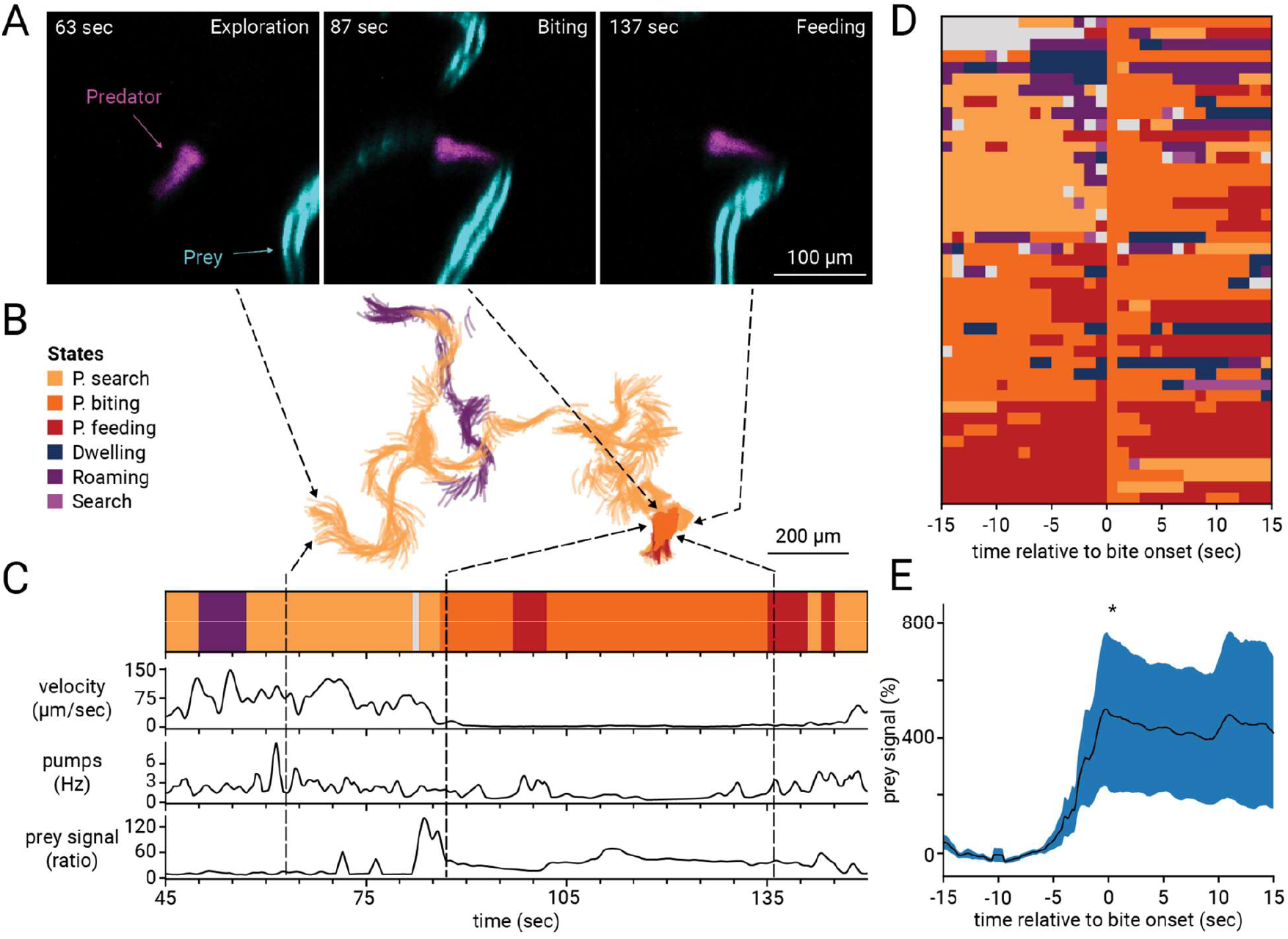
Visualizing animal and environment interaction. (A) Example images of sequential stages of a predatory event involving the predator *P. pacificus* expressing RFP in the pharynx (magenta) and *C. elegans* prey expressing GFP in the body wall muscles (cyan) at 3.1x magnification. Scale bar corresponds to 100 µm. Arrows point to the corresponding track in B. (B) Example tracks of the centerline of the predator between 45 sec and 150 sec of recording, colors indicate predicted behavioral states using the model published in. Scale bar corresponds to 200 µm. (C) Ethogram showing the predicted behavioral states over time, as well as velocity, pumping rate, and prey signal within a 34 µm circular mask centered at the anterior end of the predator. Arrows point to the corresponding track in B. (D) Stacked ethograms of n=43 tracks, N=8 predators aligned at the onset of predatory biting at t=0 (E) Prey signal (%) of the tracks shown in D (mean ± s.e.m.), excluding those with the mode of predatory biting or predatory feeding, n=17, N=7. Prey signal was detected as in C, but normalised to the signal within -15 to -5 sec before bite onset at t=0. Statistics performed with an upper-tailed paired t-test comparing time ranges -15 to -5 and 0 to 15, * p < 0.05.

A section of the recording is displayed in Fig. 4B-C, showing the predator’s centerline and corresponding metrics: velocity and pumping rate. Tracking *P. pacificus* on prey revealed that during biting and feeding states, velocity decreased while pumping rates increased; in contrast, exploration featured higher velocity and lower pumping rates. By aligning multiple ethograms to biting events (Fig. 4D), we could show that GFP prey signal significantly increased at bite onset (Fig. 4E), reflecting predator-prey contact and ingestion of the green prey. These measurements allowed us to verify the model predictions and confirm that biting states occurred in the presence and close contact with prey. The microscope is therefore suitable for imaging small, moving animals within their con-specifics or other relevant species, thus enhancing our ability to understand multi-species interactions.

### Long-term tracking of organ activity in behaving animals using the single-color microscope

Thanks to its modular design, the microscope also supports single-color imaging with a simplified optical path (Figure S2), offering twice the FOV for a single fluorescent label or dye. The single-color microscope uses a subset of components from the dual-color microscope as it does not require the image splitter (Fig. 5A, B). Similar to the dual-color setup, the magnification can be adjusted to suit specific applications. For instance, we used it here to image the feeding organ of *C. elegans* (Fig. 5C). With the same objectives and magnifications as the dual-color microscope, the design uses relatively low effective magnification and low NA, resulting in a large depth of field. This feature is particularly advantageous as it minimizes motion artifacts and is ideal for imaging entire animals or tracking features within freely moving animals, such as cells or organs that may move out of the focal plane.

**Figure 5:**
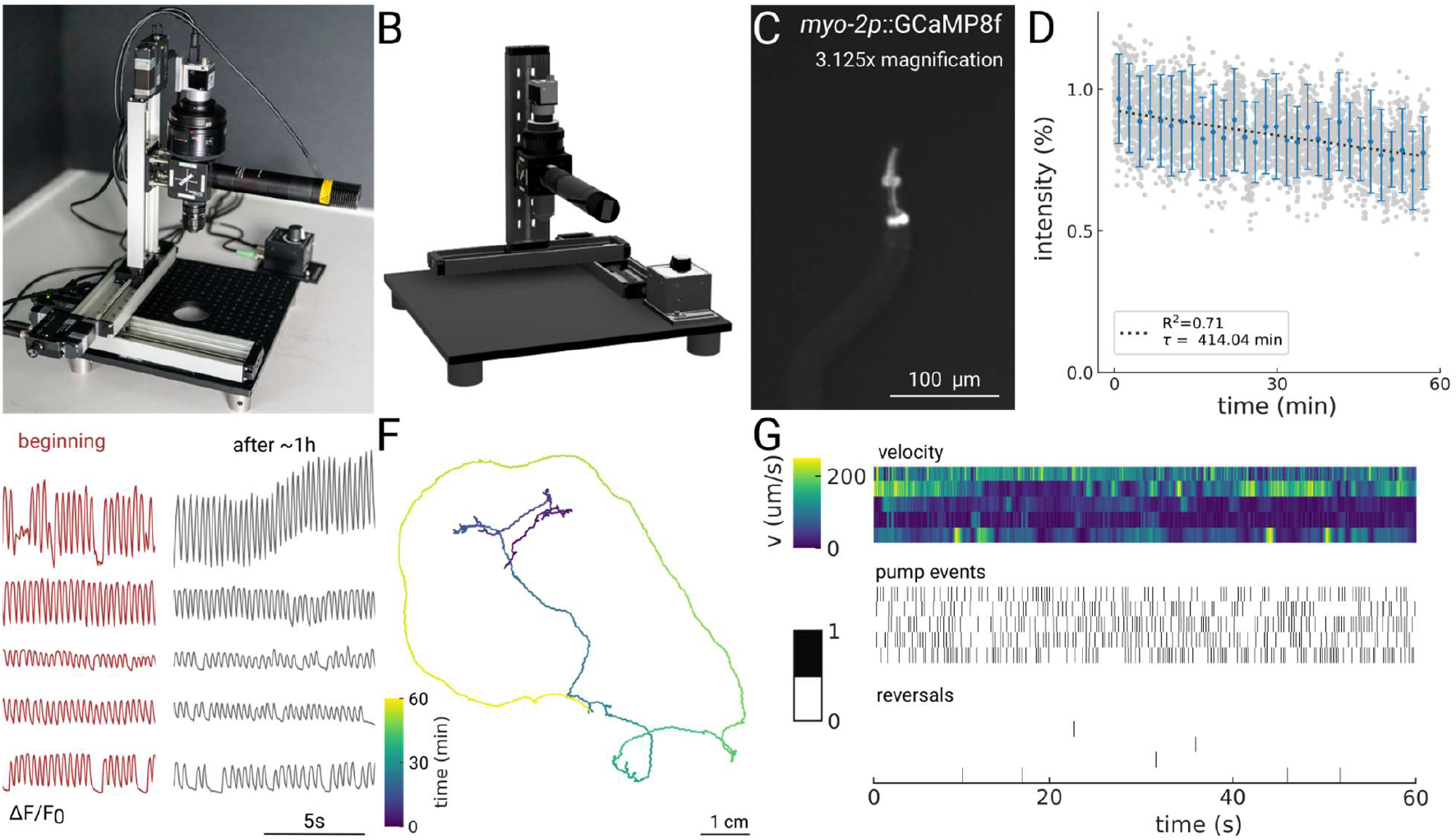
Modular single-color design enables long-term tracking. (A) Setup adapted for single color imaging. This version uses a subset of parts from the dual-color microscope. (B) 3D rendered model of the setup. (C) Example image of a *C. elegans* nematode with the pharyngeal muscle expressing GCaMP8f. (D) Photobleaching curves for 1h of continuous tracking and illumination at 2.78x. (E) Example signals from muscle contractions measured from the GCaMP8f indicator at the beginning and end of the 1h track for N = 5 animals. (F) Example track of a worm. (G) Ethograms for N=5 animals for 1 minute of data showing speed, feeding, and reversals.

To evaluate whether fluorescence levels and tracking stability are suitable for hour-long experiments, we designed an experiment tracking a single *C. elegans* nematode while foraging. We use the microscope to automatically follow the nematodes with the pharynx expressing a genetically-encoded calcium indicator^40^. We find that over the course of 1 h recordings, the mean intensity is only reduced by ∼ 25%, indicating low photobleaching (Fig. 5D). The resulting signals from the calcium indicator are still sufficient to extract peaks, corresponding to contractions of the feeding muscle after ∼ 1h of recording (Fig. 5E). The tracking stage can automatically retain the animals in the field-of-view while they travel over multiple centimeters (Fig. 5F), allowing us to observe unrestrained behaviors. This configuration allows simultaneous measurement of multiple behaviors, such as feeding, reversals, and locomotion speed (Fig. 5G). During long-term experiments, the animals remained viable with no significant changes in velocity or pumping rate observed, suggesting that the experimental conditions did not adversely affect the viability of the animals over time. This stability in behavior enables investigations into internal states, development, or sleep^47^.

### Brightfield tracking for non-model species

Brightfield imaging is versatile and amenable for species in which the ability to use transgenics for labeling body parts is not established. To demonstrate the capability of the microscope for brightfield tracking, we recorded locomotion behavior in a non-model species, namely the tardigrade *H. exemplaris* (Fig. 6A). By training the pose estimation classifier DeepLabCut, we could predict the position of all eight legs, eyes, as well as the most anterior and posterior ends of the body (Fig. 6B-C). The model performed well, with a mean prediction error of 6.2 pixels (3.53 µm) on the test set and 1 pixel (0.57 µm) on the training set. However, accuracy was lower in occluded or out-of-focus areas (not shown).

**Figure 6:**
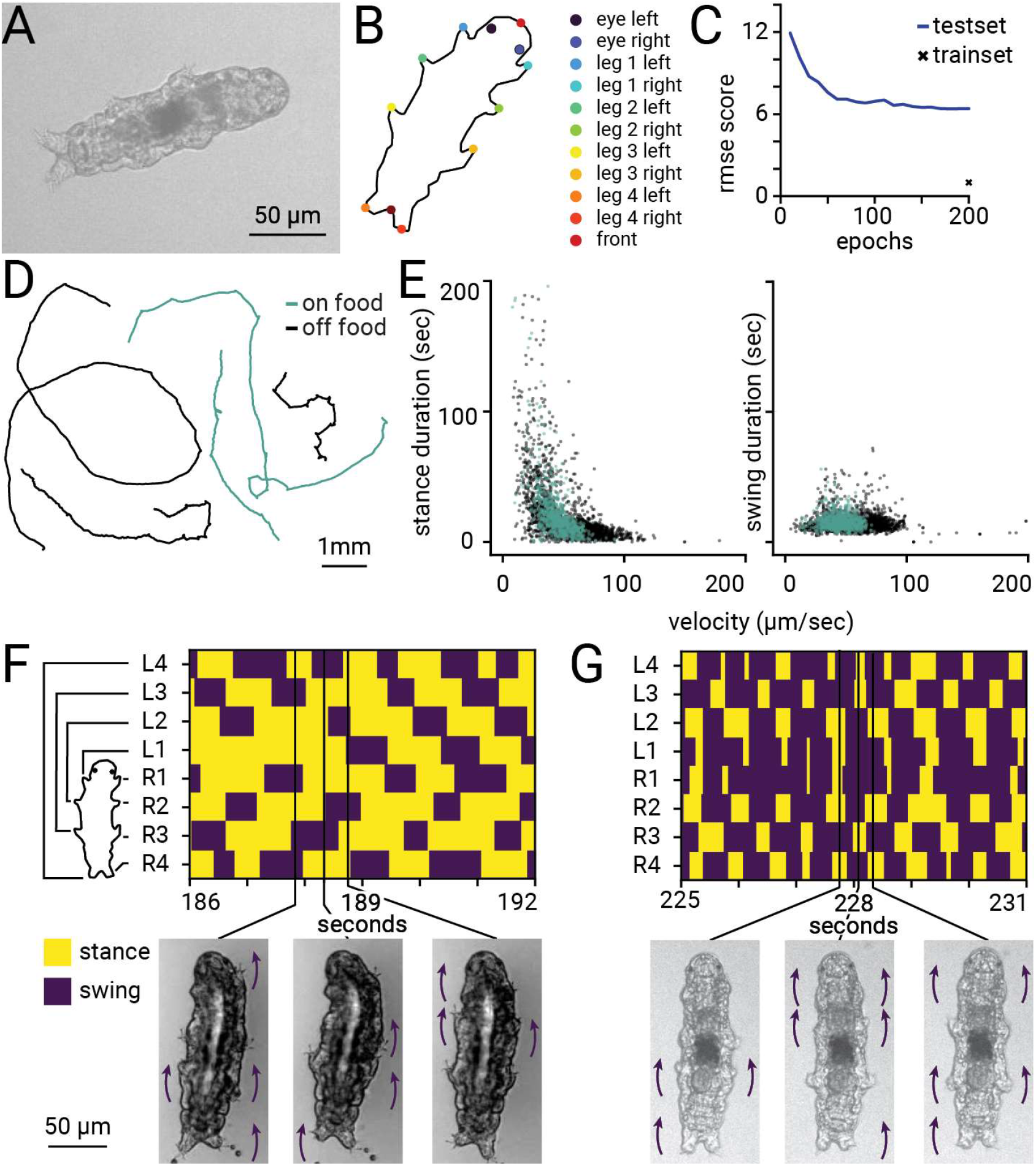
Brightfield tracking for non-model species. (A) Example image of a tardigrade on the microscope. (B) Labels used for pose estimation with DeepLabCut (C) DeepLabCut pixel error across training epochs for the test set and on the epoch 200 for the training set (D) Tracks of tardigrades (E) Speed distribution of N=5 tardigrades on (green) and off food (black), only subparts of videos analysed where tardigrades where visible and in focus. (F) example tetrapod gait (G) example tetrapod gallop gait.

We tracked animals in both the presence and absence of their algal food source, but the tracks did not show notable differences (Fig. 6D). As observed in other species, such as insects ^48–50^, we found that in *H. exemplaris*, while the duration of the stance decreases with the speed of the animal, the swing duration remains constant across speed changes (Fig. 6E). As previously described^51,52^, we recover two prominent recurring gait types, both shown in Fig. 6F-G. In the first, the tetrapod gait (Fig. 6F), tardigrades move their legs sequentially, with the stance of the next posterior leg following that of the next anterior leg. The left and right sides alternate, with one side following shortly after the other. One gait cycle seems to overlap with the following one. In contrast, during the second gait type, termed tetrapod gallop ^52^, the two leg pairs 2 and 3 are sequentially moved, with the left and right pairs moving simultaneously (Fig. 6G). However, the two gait types appear to continuously transition into one another. For example, at 186 sec in Fig. 6F, the left and right legs of pair 2 move simultaneously. By imaging animals navigating over large distances, we capture more strides per animal than previous studies, likely enabling the observation of gait transitions. Further studies could investigate how gait choice depends on environmental parameters such as substrate stiffness ^51^, or how it supports different navigational goals.

## Discussion

We presented a tracking epifluorescence microscope using only commercially available parts, paired with an integrated and fully automated software application. Currently, similar microscopes are built in laboratories by experts in optical design and hardware control, or expensive commercial solutions have to be found, which are often less adaptable to diverse experiments. Our goal was to create a microscope accessible to users with less experience in building optical setups, yet sufficient for producing research-quality data. While some parts could be replaced by hardware made in a mechanical or electrical workshop, we purposely decided to suggest only commercially available components to make the design accessible to novice users or users without access to such facilities. Our website suggests which parts could be replaced by machining or laser cutting, and potential up- or downgrades of components depending on the specific use case.

We purposely selected lower magnifications than commonly used for calcium imaging in microscale animals, as it allowed for a wider range of applications and organisms. Specialized hardware, for example, the ‘Wormspy’ system, excels in high-resolution imaging of cellular processes^53^, while our setup offers flexibility and accessibility for diverse experimental setups. Both systems address different needs in the field, with Wormspy ideal for detailed cellular studies and our microscope suitable for broader behavioral and ecological investigations. Moreover, our setup could be adapted by using a tube lens with a longer focal distance to achieve higher magnification, suitable for neuronal imaging of smaller cells.

Our microscope is versatile, allowing tracking and measurement of functional properties of muscles and neurons, as demonstrated in larval Drosophila crawling, *C. elegans* locomotion, and touch receptor neuron activity. The dual-color feature unlocks possibilities for studying multi-animal and multi-species interactions, bridging the gap between ethology and ecology. For instance, it could shed light on intriguing behaviors like stress-induced cannibalism in Drosophila larvae ^54^. Similarly, the microscope could be used to track individuals within collectives, by labeling a single or a few animals, relevant to investigate collective behaviors^55,56^.

While the microscope is capable of showing neuronal activity under specific conditions, it is optimized for fast, large field-of-view imaging rather than sensitivity for small neuronal signals. However, only specialized hardware solutions for ratiometric neuronal imaging in freely moving animals exist for *C. elegans*, Drosophila larvae, and also zebrafish. We expect that the proposed microscope fills a niche between these more involved setups and the easier behavioral setups using brightfield microscopes.

## Methods

### C. elegans maintenance

All nematode strains were cultured on Nematode Growth Media (NGM) plates with OP50 at 20°C.

| Species | Designation | Identifiers | Additional information | Source or reference |
| --- | --- | --- | --- | --- |
| *C. elegans* | *zcls14 [myo-3::GFP(mit)]* | SJ4103 |  | CGC |
| *C. elegans* | *nonEx106[myo-2p::GCaMP8f]* | INF418 | Integrated from INF418 | This publication |
| *D. melanogaster* | *MEF2-Gal4, UAS-mGerry* | MEF2-Gal4 |  | Heckscher lab, University of Chicago |
| *P. pacificus* | *Ppa-myo-2p::RFP* | JWL27 |  | 44 |
| *C. elegans* | *lite-1(ce314) gur-3(ok2245) X; nonEx133[Pmec-17_GCaMP8f_SL2_mKate2_let-858_3'UTR + Punc-122::tagBFP]* | INF204 |  | 19 |
| *H. exemplaris* |  |  |  | Gift from Jean-Paul Concordet, Muséum National d'Histoire Naturelle, France |

### Imaging Plates

All experiments were performed on imaging NGM plates. In contrast to NGM plates, imaging plates contain 17 g of agarose instead of agar and no cholesterol for 1 L medium.

### Recordings

For all experiments, the GlowTracker software was used to record images and stage position during tracking.

### Long-term tracking Assay

Long-term assays were performed on 10 cm imaging NGM plates containing a concentrated patch of fresh OP50 (10X concentrated from overnight culture). *C. elegans* young adults were selected and starved for 2 h before being transferred individually to the NGM plate. Following a 15-minute acclimation period under blue light, the worm locomotion was recorded for 1h at 3.125x magnification using the single-color configuration with a 15 ms exposure and a sampling rate of 30 frames/s.

### Analysis of long-term C. elegans foraging data

The center-of-mass coordinates as well as intensity-related measures such as mean intensity, maximum intensity, and centerline of the *C. elegans* pharynx were extracted using PharaGlow ^24^. Using a peak-detection algorithm, intensity peaks of the GCaMP8f indicator were annotated and used to calculate a pumping rate, similar to the process described in ^24,40^.

### Drosophila maintenance and strains

All fly stocks were maintained at 25°C. Flies expressing MEF2-Gal4 (Expresses GAL4 in muscle cells, likely BDSC_27390), UAS-mGerry (GCaMP6m fused to mCherry under the control of UAS, BDSC_80141) were provided by E. Heckscher. The genotype of the flies in the experiments was w*, MEF2-Gal4, UAS-mGerry.

### Single Drosophila larvae chemotaxis Assay

All chemotaxis assays involved early and late second-instar larvae tested during the day. Room temperature was maintained at around 20□, and relative humidity of 30 ± 6 %. Chemotaxis assays were performed on 15 cm unseeded imaging NGM plates. At one end of the plate, 20 μL of undiluted commercial apple vinegar (REWE, Germany) was deposited. A single larva was positioned within a droplet of distilled water approximately 3 cm from the vinegar drop. Their locomotor movement was videotaped for 5 min at 1x magnification with 20 ms exposure using the dual-color configuration and a sampling rate of 19.6 frames per second for 5 minutes.

### *Drosophila* crawling analysis

Dual-color images were analyzed with PharaGlow ^24^. The segmentation was adapted to reliably detect the larvae (Fig. S2F), virtually straighten the animal, and extract kymographs of intensity along the anterior-posterior axis. The red channel R(t) was used to correct motion artifacts of the calcium signal (G(t)) using the same approach as described in ^14^. In brief, for each location on the anterior-posterior axis (L), we estimate α(L), such that the corrected signal is F_corr =(G−αR)−⟨G−αR⟩, with α(L) minimizing ∑(G(t)−αR(t))^2^.

### Calculating resulting magnification

For fixed focal distance objectives (e.g., 50 mm Yongnuo, 12 mm EO), the magnification is given by *M* = *f*_*TL*_/*f*_*OBJ*_ .As the microscope uses a non-standard tube lens (*f*_*TL, GlowTracker*_= 50 mm), the magnification of the attached objectives changes from the nominally stated magnification. For commercial objectives such as the Olympus 10x and Olympus 20x, the magnification can be calculated as 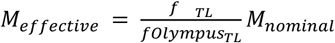. For an Olympus 10x objective with *f*_*TL*_,^*Olympus*^ = 180 *mm, f*_*TL*_= 50 *mm*, and *M*_*nominal*_= 10*x*, this results in *M*_*effective*=_ 50/180 ∗ 10 ≃ 2.78.

### Predation Assay

Young *C. elegans* larvae (SJ4103) were washed with M9, passed through a 20 µm filter twice, centrifuged, and deposited on an unseeded 10 cm NGM imaging plate by pipetting 4 µL of the worm pellet. The plate was left undisturbed for at least 1 hour to allow the larvae to evenly distribute on the plate. Following a 2-hour starvation period, six washed young adult *Pristionchus pacificus* (JWL27) predatory hermaphrodites were introduced to the plate. Animals were imaged with a 16 mm objective at 30 fps using the dual-color microscope. Two neutral density filters with an optical density (OD) of 0.6 and 1.0 were added to reduce the intensity of the red imaging channel, as the fluorescence of *Ppa-myo-2p::tagRFP* of the adult predator was stronger than the label of the prey, making the adjustment of the dynamical range of the camera difficult for both colors simultaneously.

### Statistical Analysis of Prey Signal

The recordings were analysed with a custom Python script, incorporating the imaging analysis tool PharaGlow^24^, to extract the pharynx centerline, center of mass, and additional metrics of the predator (*P. pacificus*). The coordinates of the recordings were transformed into the stage space and used for velocity calculation. Following this, the behavior of the predators were analysed with a machine-learning based model published to detect behavioral states^44^. To analyse behavior and prey signal around the onset of predatory biting, the tracks were aligned at the onset of predatory biting events. Further, the extracted centerline was used to measure the prey signal in the green channel. For this, a 34 µm wide circular mask, centered around the anterior end of the centerline was used. The prey signal was calculated from the raw GFP signal as the ratio: *preysignal* = (*GFP*_95*percentile*_ − *GFP*_5 *percentile*_)/*GFP*_5 *percentile*_ To analyse the rise of prey signal at biting onset, the prey signal of each track was normalised to a baseline defined as the mean of prey signal between 15 to 5 sec before bite onset, as *preysignal* (%) = (*preysignal* − *baseline*)/*baseline*. Statistics were performed using a paired t-test, comparing the time ranges of 15 to 5 sec before bite onset and 0 to 15 sec.

### Calcium Imaging of PLM

*C. elegans* young adults expressing *mec-17p::GCaMP8f::mCherry2* were transferred 15 min before recording to an NGM imaging plate, and shortly before the recording, the agar was cut and transferred to the recording setup. The NGM plates were seeded with 50 μL of OP50 the evening before the experiment and allowed to grow overnight. Animals were recorded in a dual-color microscope, as described in Table S2. A tube lens of type Yongnuo 50 mm and an objective of type EO 12 mm were used, resulting in an effective magnification of 4.1x (Table S1). The red channel was used for tracking the animal, and the intensity of the red channel was attenuated using an OD=1 neutral density filter.

### Touch stimulation during calcium imaging

Animals were stimulated with a sinusoidal wave at 630 Hz and 20 Vpp using a piezo buzzer as described in ^40^. The stimulation protocol had five repeats of 1 sec with an interstimulus interval of 30 s and a pre-stimulation period of 30 s. The stimulation was controlled via a custom MATLAB script and an NI DAQ board (NI, TX, USA).

Extraction of the calcium signals was performed with custom Python scripts. In brief, after cropping the images to a 50×50 px region of interest, the background (defined as the 50th percentile of pixel intensity in every 100th frame) was subtracted from the red channel. Neurons were detected by blurring with a Gaussian filter (st. dev = 0.3) and the neuron mask was found using thresholding. The mask was then used to extract the raw signal from the reference and signal images. For ratiometric analysis, dR/R0 was calculated as

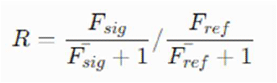

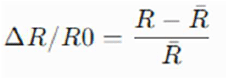

Velocity was calculated directly from stage coordinates, which were recorded at 30 fps.

### Statistical analysis of calcium imaging

For plotting and statistical analysis, the traces were aligned at the stimulus onset. For the statistical analysis, we used a dependent t-test for paired samples to compare the mean value of the prestimulus interval (from -10 sec to 0 sec aligned to the stimulus) and the stimulus interval (0 sec to 1 sec) within one stimulus repeat. For figures 3E and F, the repeats were filtered according to reversal events occurring within 2 sec of stimulus onset. Reversals were defined based on the method described in ^57^ by detecting sharp angular changes in the trajectory.

The effect of reversals on the mean post-stimulus calcium signal was tested using a mixed linear model with PLM activity as the dependent variable and reversals as the independent variable, categorical, while accounting for random effects between worms. The analysis was performed with the ‘statsmodels’ package in Python.

### Tardigrade maintenance

Tardigrades were kept at room temperature (20°C) in Volvic water with a suspension of Chlorella vulgaris grown in BG11 medium added regularly as a food source. Animals of approximately the same size were selected from a culture plate for tracking.

### Brightfield tracking of non-model species

Tardigrades were observed on 15 cm unseeded imaging NGM plates containing Volvic water. Locomotion was recorded using a 12 mm EO objective at 4.1x magnification in bright-field mode, employing a single-color configuration with red LED illumination (650 nm) and a 50/50 image splitter instead of a dichroic mirror. Images were captured at 30 frames per second with an exposure of 96 μs. The tardigrades’ center of mass was tracked in the X-Y plane for 3-5 minutes using the GlowTracker software.

The pose estimation was done with DeepLabCut (3.0.0rc4). We used nine videos for training, three on food (algae), and six off food. From those, we extracted and labeled 179 frames. Labeling was done as shown in Fig. 6B: we labeled the eyes (left and right), all four leg pairs (each left and right), and two additional points at the anterior and posterior ends of the body.

For training, we used 80% of the labeled data. We trained a ResNet-50-based neural network with default parameters for 200 training epochs. This was sufficient to achieve a test error of 6.41 pixels, while the training error was 1 pixel.

For prediction and analysis, a subset of videos was used for training. This was because some videos had a large fraction of frames where the tardigrades were either not in focus, in an undesired position, or obscured by algae. Those videos were included in the training to make the model robust to perturbations. Furthermore, only those parts of the recordings were used for analysis, where the tardigrades were clearly visible and moving forward (except for Fig. 6C, where the full tracks are shown).

For analysis, video recordings were aligned with the stage coordinates. The center of mass was calculated as the mean of all tracked body parts. For each leg, its velocity was calculated. Swing bouts were extracted by peak detection, including the width of the peak, using scipy.find_peaks. The mode was calculated for bouts that were shorter than five frames (at 30 fps). Stances were identified as the invert of swings.

### 3D model of the Microscope

The 3D model of the single and dual-color microscope was created using the Fusion 360 software under the license provided by the Max Planck Digital Library.

## Data and code availability statement

The Glowtracker software is available on GitHub https://github.com/scholz-lab/GlowTracker and installable via the Python package manager pip as “glowtracker”. Further documentation and instructions are available at https://scholz-lab.github.io/GlowTracker.

## Acknowledgements

We thank Ellie Heckscher and Matthieu Louis for reagents. We thank Louis Wolinski for assistance with the 3D models. We thank members of the laboratory of James Lightfoot (Fumie Hiramatsu and Desiree Goetting) for testing our documentation by building a microscope and providing valuable feedback.

