## Supplementary for "A modular multi-color fluorescence microscope for simultaneous tracking of cellular activity and behavior"

Supplementary Information

**Benchmarking of the live analysis and tracking performance**

GlowTracker performance is limited by two aspects: Image Acquisition and Tracking. Image Acquisition refers to the acquisition and processing of images from the camera until receiving a ready-to-use image in the host machine, and Tracking refers to the computation of key point positions in the image and stage adjustments. Several factors influence the Image Acquisition such as the exposure time which determines how long the sensor is exposed to light before being read out, the image size, and the binning mode. Shorter exposure times and smaller image sizes result in higher acquisition rates. Binning modes, like additive modes, can increase the brightness but reduce the effective image resolution. Therefore, personalizing these factors depends on the experimental setup and the organism under study. To optimize the Image Acquisition, GlowTracker employs a rolling shutter mode, consecutively exposing sensor rows with minimal time offsets (8 µs in our model). This reduces sensor readout wait time, improving the effective acquisition rate. Following successful image acquisition, the application engages in Tracking by computing the location of interest, adjusting the stage accordingly, and awaiting the next image for tracking.

To determine where the tracked animal moved requires calculating the offset of the most-likely object from the center. To this end, multiple image denoising and thresholding steps are used to detect the largest object closest to the center of the image. These coordinates are then handed to the tracking algorithm which calculates the compensatory movement of the stage in real-world coordinates required to re-center the animal. To speed up the calculation, users can determine a resizing parameter (Fig. S1A) which will determine the downsampling applied to images before analysis. To maintain accuracy of tracking, we find that going beyond a resize factor of 6 does not meaningfully improve image analysis speeds, while reducing the accuracy of the calculated tracking correction.

Benchmarking is performed with maximum image ROI (3088 x 2064 pixels), no binning, in a laptop with 12th Gen Intel(R) Core(TM) i7-1255U 1.70 GHz CPU, 16 GB of RAM, and on a Windows 10 64-bit operating system. The evaluation focuses on the effective image-tracking time reflecting the timestamp at the initiation of tracking. The relationship between the effective image acquisition rate and the effective tracking rate is nearly linear, with faster image acquisition leading to quicker tracking (Fig. S1B). Calculating the Frames per track ratio (image acquisition rate/tracking rate) provides insight into the number of frames acquired to complete the tracking of one frame. The ratio is mostly linearly proportional to the image acquisition rate because the time it takes to compute tracking is fixed, regardless of how quickly images are acquired (Fig. S1A).

The performance of GlowTracker varies based on hardware, software, and the subject being studied. In experiments with *Caenorhabditis elegans* and *Pristionchus pacificus*, we identified an effective exposure time range between 20 ms to 60 ms for obtaining high-quality images while maintaining optimal responsiveness of the stage. This range yields acquisition rates from 50 Hz to 16.67 Hz and tracking rates from 11 Hz to 6.5 Hz, resulting in frames-per-track ratios ranging from 5 to 2.5 times.

**Upgrade options**

There are several options to adapt the microscope for advanced experimental needs. Arena sizes can be expanded by substituting stages with longer ranges. The camera can be substituted by any similar camera model using USB3.0 to fit the needs best, for example, cameras with slower but more sensitive chips. However, we found that the model used here is suitably fast and has sufficient sensitivity (80% quantum efficiency peak) to image samples with low expression of genetically-encoded indicators.

**Supplementary table 1: Objectives and magnifications**

Magnification and expected field-of-view for objectives used with the GlowTracker. The numerical aperture (NA) is given as reported by the manufacturer, and therefore reflects the maximum possible NA. Asterisks indicate the objective was tested for compatibility as an upgrade option, but is not used in this paper.

| **Objective** | **f#** | **NA =½*f#** | **Focal length (mm)** | **Max FOV(mm)** | **Magnification** |
| --- | --- | --- | --- | --- | --- |
| **Yongnuo 50 mm** | f1.8 - f22 | 0.27 | 50 | 7.41 x 4.95 | 1 |
| **EO 16 mm** | f/1.6 - f/16 | 0.31 | 16 | 2.3 x 1.59 | 3.1 |
| **EO 12 mm** | f/1.8 - f/16 | 0.22 | 12 | 1.8 x 1.2 | 4.1 |
| **Olympus 10x*** | f/2 | 0.25 | 18 | 2.74 x 1.83 | 2.7 |
| **Olympus 20x*** | f/1.25 | 0.40 | 9 | 1.32 x 0.88 | 5.6 |

**Supplementary table 2. Components for the dual color microscope**

|  |  | **Part** | **Description** | **Pc** | **Price per unit € (08/2024)** | **Vendor** | **Link** |
| --- | --- | --- | --- | --- | --- | --- | --- |
| **Lightpath** | 1 | Camera Cable | USB 3.0 Kabel Stecker A an Stecker Micro B, Premium, AWG28 / AWG24, UL, KUPFER, schwarz, 1m | 1 | 7,90 | Kabelmeister | https://www.kabelmeister.de/USB-3.0-Kabel-Stecker-A-an-Stecker-Micro-B-Premium-AWG28-AWG24-UL-KUPFER-schwarz-1m/UK30P-AMB-010S |
|  | 2 | Camera | Basler ace U  acA3088-57um | 1 | 419 | Blaser | https://www.baslerweb.com/en/shop/aca3088-57um/ |
|  | 3 | Kipon Canon EOS to C-Mount | C-Mount to Canon EOS EF/EF-S Lens Mount Adapter | 1 | 59 | [Kipon](https://kipon.de/shop/adapter/mechanische-adapter/canon-eos-c/) or [B&H](https://www.bhphotovideo.com/c/product/1458022-REG/kipon_canon_eos_c_canon_eos_to_c.html) | https://www.bhphotovideo.com/c/product/1458022-REG/kipon_canon_eos_c_canon_eos_to_c.html |
|  | 4 | 50mm objective | YN50mm F1.8: 50mm F1.8 ObjectiveI | 2 | 110 | YONGNUO | https://th.hkyongnuo.com/products/yn50mm-f18 |
|  | 5 | SM1A2: Adapter | External SM1 Threads and Internal SM2 Threads | 2 | 25,82 | Thorlabs | https://www.thorlabs.com/thorproduct.cfm?partnumber=SM1A2 |
|  | 6 | Filter Cube | DFM1/M: Kinematic Fluorescence Filter Cube for Ø25 mm Fluorescence Filters, 30 mm Cage Compatible | 3 | 385,34 | Thorlabs | https://www.thorlabs.com/thorproduct.cfm?partnumber=DFM1/M |
|  | 7 | Threaded End Cap for Machining | SM1CP2M: Externally SM1 | 3 | 19,09 | Thorlabs | https://www.thorlabs.de/thorproduct.cfm?partnumber=SM1CP2M |
|  | 8 | Cage Assembly Rod | ER025: Cage Assembly Rod, 1/4” Long, Ø6 mm | 16 | 4,96 | Thorlabs | https://www.thorlabs.de/thorproduct.cfm?partnumber=ER025 |
|  | 9 | KCB1EC/M : Elliptical Mirror Bores | Right-Angle Kinematic Elliptical Mirror Mount with Smooth Cage Rod Bores | 2 | 202,55 | Thorlabs | https://www.thorlabs.com/thorproduct.cfm?partnumber=KCB1EC/M#ad-image-1 |
|  | 10 | Elliptical Mirror | BBE1-E02: 1” Broadband Dielectric, 400 - 750 nm | 1 | 118,53 | Thorlabs | https://www.thorlabs.de/thorproduct.cfm?partnumber=BBE1-E02 |
|  | 11 | C4W-CC: 30 mm Cage Cube Connector | 30 mm Cage Cube Connector for C4W and C6WR Series Cubes | 1 | 54,54 | Thorlabs | https://www.thorlabs.com/thorproduct.cfm?partnumber=C4W-CC#ad-image-0 |
|  | 12 | External Threads | SM1 (1.035"-40) Coupler, External Threads, 0.25" Long | 1 | 19,96 | Thorlabs | https://www.thorlabs.de/thorproduct.cfm?partnumber=SM1T3 |
|  | 13 | AC254-050-A-ML | f=50 mm, Ø1” Achromatic Doublet, SM1-Threaded Mount, ARC: 400-700 nm | 1 | 105,83 | Thorlabs | https://www.thorlabs.de/thorproduct.cfm?partnumber=AC254-050-A-ML |
|  | 14 | Lens Tube Without External Threads | SM1M05 1/2” Long, Two Retaining Rings Included | 1 | 13,96 | Thorlabs | https://www.thorlabs.de/thorproduct.cfm?partnumber=SM1M05 |
|  | 15 | Zoom Housing for Ø1” Optics | SM1NR1, Non-Rotating, 2” travel | 1 | 214,89 | Thorlabs | https://www.thorlabs.de/thorproduct.cfm?partnumber=SM1NR1 |
|  | 16 | Precision slit CDI11786 | 3500um ± 40um width. 10 mm length. 316 Stainless Steel | 1 | 142,71 | TBD from Thorlabs | request CDI11786 from Thorlabs |
|  | 17 | Lens Tube Spacer | SM1S20: Lens Tube Spacer, 2” Long | 1 | 15,44 | Thorlabs | https://www.thorlabs.de/thorproduct.cfm?partnumber=SM1S20 |
|  | 18 | ACL25416U-A - Aspheric Condenser | Lens, Ø1”, f=16 mm, NA=0.79, ARC: 350-700 nm | 1 | 29,16 | Thorlabs | https://www.thorlabs.de/thorproduct.cfm?partnumber=ACL25416U-A |
|  | 19 | SM1 Lens Tube | SM1L10 : 1.00" Thread Depth, One Retaining Ring Included | 1 | 14,01 | Thorlabs | https://www.thorlabs.de/thorproduct.cfm?partnumber=SM1L10 |
|  | 20 | Rod Adapter | ERSCB-P4: Rod Adapter for Ø6 mm ER Rods, L = 0.27”, 4 Pack | 1 | 58,41 | Thorlabs | https://www.thorlabs.com/thorproduct.cfm?partnumber=ERSCB-P4 |
|  | 21 | Cage Assembly Rod | ER025: Cage Assembly Rod, 1/4” Long, Ø6 mm | 4 | 4.96 | Thorlabs | https://www.thorlabs.de/thorproduct.cfm?partnumber=ER025 |
|  | 22 | SM2A53: Adapter with External | M52 x 0.75 Threads and Internal SM2 Threads | 2 | 21,49 | Thorlabs | https://www.thorlabs.de/thorproduct.cfm?partnumber=SM2A53 |
|  | 23 | SM1RR: SM1 Retaining Ring Ø1” | SM1 Retaining Ring for Ø1" Lens Tubes and Mounts | 4 | 4,43 | Thorlabs | https://www.thorlabs.com/thorproduct.cfm?partnumber=SM1RR |
| **Filters** | 24 | Red Emission filter | FF01-618/50-25: 618/50 BrightLine® single-band bandpass filter | 1 | 405 | Semrock | https://www.idex-hs.com/store/product-detail/ff01_618_50_25/fl-004072 |
|  | 25 | Green Emission filter | 67-030: 520nm CWL, 25mm Dia, 36nm Bandwidth, OD 6 Fluorescence Filter | 1 | 313 | Edmund Optics | https://www.edmundoptics.eu/p/520nm-cwl-25mm-dia-36nm-bandwidth-od-6-fluorescence-filter/21570// |
|  | 26 | Excitation Filter | 59011x: Dual-band-pass Excitation Filter | 1 | 400 | Chroma | https://www.chroma.com/products/parts/59011x |
|  | 27 | Dichroic Beamsplitter  562 nm | FF562-Di03-25x36: 562 nm edge BrightLine® single-edge standard epi-fluorescence dichroic beamsplitter | 2 | 305 | Semrock | https://www.idex-hs.com/store/product-detail/ff562_di03_25x36/fl-007122 |
|  | 28 | Dichroic Beamsplitter  488/561 nm | Di01-R488/561-25x36: 488/561 nm lasers BrightLine® dual-edge laser dichroic beamsplitter | 1 | 555 | Semrock | https://www.idex-hs.com/store/product-detail/di01_r488_561_25x36/fl-006697 |
| **ND Filters** | 29 | ND Filters | ND filter OD = 1.0 and OD = 0.5 | 1 | 673.81 | Thorlabs | https://www.thorlabs.de/thorproduct.cfm?partnumber=NDK01 |
| **Illumi-**  **nation** | 30 | MNWHL4: White LED | 4900 K, 740 mW(Min) Mounted LED,1225 mA | 1 | 158,81 | Thorlabs | https://www.thorlabs.de/thorproduct.cfm?partnumber=MNWHL4 |
|  | 31 | Power supply | KPS201: 15 V, 2.66 A with 3.5 mm Jack Connector | 1 | 36,66 | Thorlabs | https://www.thorlabs.de/thorproduct.cfm?partnumber=KPS201 |
|  | 32 | EDD1B: T-Cube LED Driver | 1200 mA Max Drive Current | 1 | 322,86 | Thorlabs | https://www.thorlabs.de/thorproduct.cfm?partnumber=LEDD1B |
| **Stage** | 33 | X-LSM150A: Motorized stages | Motorized linear stages with built-in controllers | 3 | 2,397 | Zaber | https://www.zaber.com/products/linear-stages/X-LSM/specs?part=X-LSM150A |
|  | 34 | Aluminum Breadboard | MSB2328/M, 230 mm x 280 mm x 9.5 mm, Ø50 mm Access Hole | 1 | 175,24 | Thorlabs | https://www.thorlabs.com/thorproduct.cfm?partnumber=MSB2328/M |
|  | 35 | Adapter Plate | AP102C : X-LRM, X-LSM, and X-LHM Metric Top Adaptor Plate | 1 | 93 | Zaber | https://www.zaber.com/products/accessories/AP102C |
|  | 36 | Bottom Adapter Plate | AP101: X-LSM Bottom Adapter Plate | 1 | 72 | Zaber | https://www.zaber.com/products/accessories/AP101 |
|  | 37 | Pillar Post | RS25/M - Ø25.0 mm Pillar Post, M6 Taps, L = 25 mm | 4 | 21.43 | Thorlabs | https://www.thorlabs.com/thorproduct.cfm?partnumber=RS25/M |
|  | 38 | AC Adapter | Power Supply, 48 V 1.25 A, Compatible with X-Series Products | 1 | 35 | Zaber | https://www.zaber.com/products/accessories/PS13S-48V12 |
|  | 39 | Data Cable | X-DC02: Data Cable, 2 ft (0.6 m), for Use with all X-Series Products | 2 | 15 | Zaber | https://www.zaber.com/products/accessories/X-DC02 |
|  | 40 | X-USBDC: USB to Serial Converter | USB to Serial Converter, with M8 Female Plug | 1 | 44 | Zaber | https://www.zaber.com/products/accessories/X-USBDC |

**Supplementary table 3.Different computers used to run the GlowTracker GUI.**

| **OS** | **CPU** | **RAM** | **Type** |
| --- | --- | --- | --- |
| Windows 10 Enterprise (64-bit) | 12th Gen Intel Core i7  1.8 GHz | 16 GB | Lenovo laptop |
| Ubuntu 18.04.6 LTS (64 bit) | 4th Gen Intel Core i7  1.9 Ghz | 8 GB | Dell Laptop |
| Windows 10 Enterprise (64-bit) | 13th Gen Intel Core i3  4.5 GHz (max) | 16 GB | Desktop (NUC) |
| MacOS (12.7.1) | Intel Core i7 2.8 GHz | 8 GB | MacBook Pro (2019) |

**Supplementary Video 1. GUI during tracking in single color**

Graphical interface of the GlowTracker App during single color tracking. The GUI allows to display the bounding box for identifying tracked objects, and toggling a tracking overlay, to display the masking done during object detection for troubleshooting.

**Supplementary Video 2. GUI during tracking in dual color**

Graphical interface of the GlowTracker App during dual color tracking. The two color channels of the image are merged for display using the estimated channel correspondence from the color calibration feature. The display frame rate is lower (15 fps) than the recording frame rate to allow usage on slower PCs.

**Supplementary Video 3. Tracking video of larvae in dual color**

Resulting images from the tracking experiment in Video 2. The scale bar is 0.5 mm. Channels were false colored in red (mCherry) and cyan (GCaMP).

**Supplementary Figures**

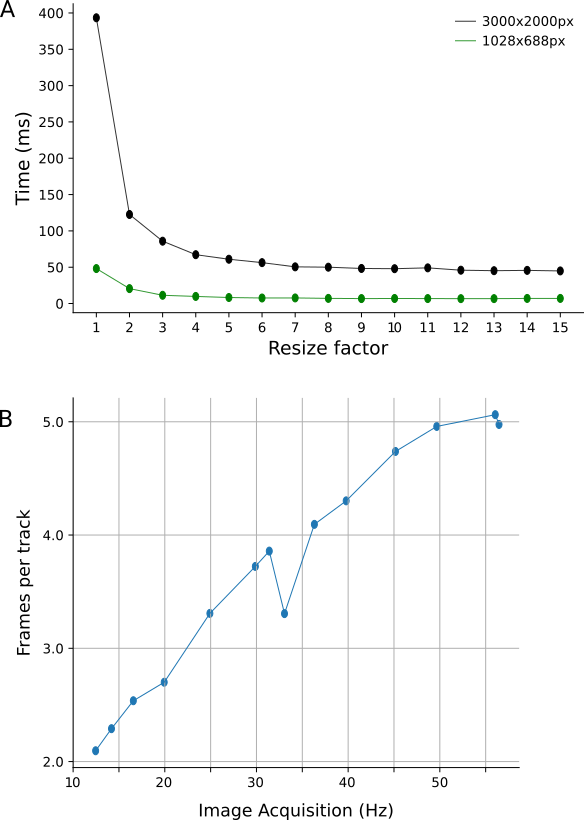

**Supplementary Figure 1. Benchmarking of GlowTracker performance**
(A) Duration of image analysis (including image denoising, animal detection, and center-of-mass calculation) for typical image resolutions. The resize factor used determines the positioning accuracy, while allowing for faster processing. (B) Tracking rate (images/track) depending on the image acquisition rate.

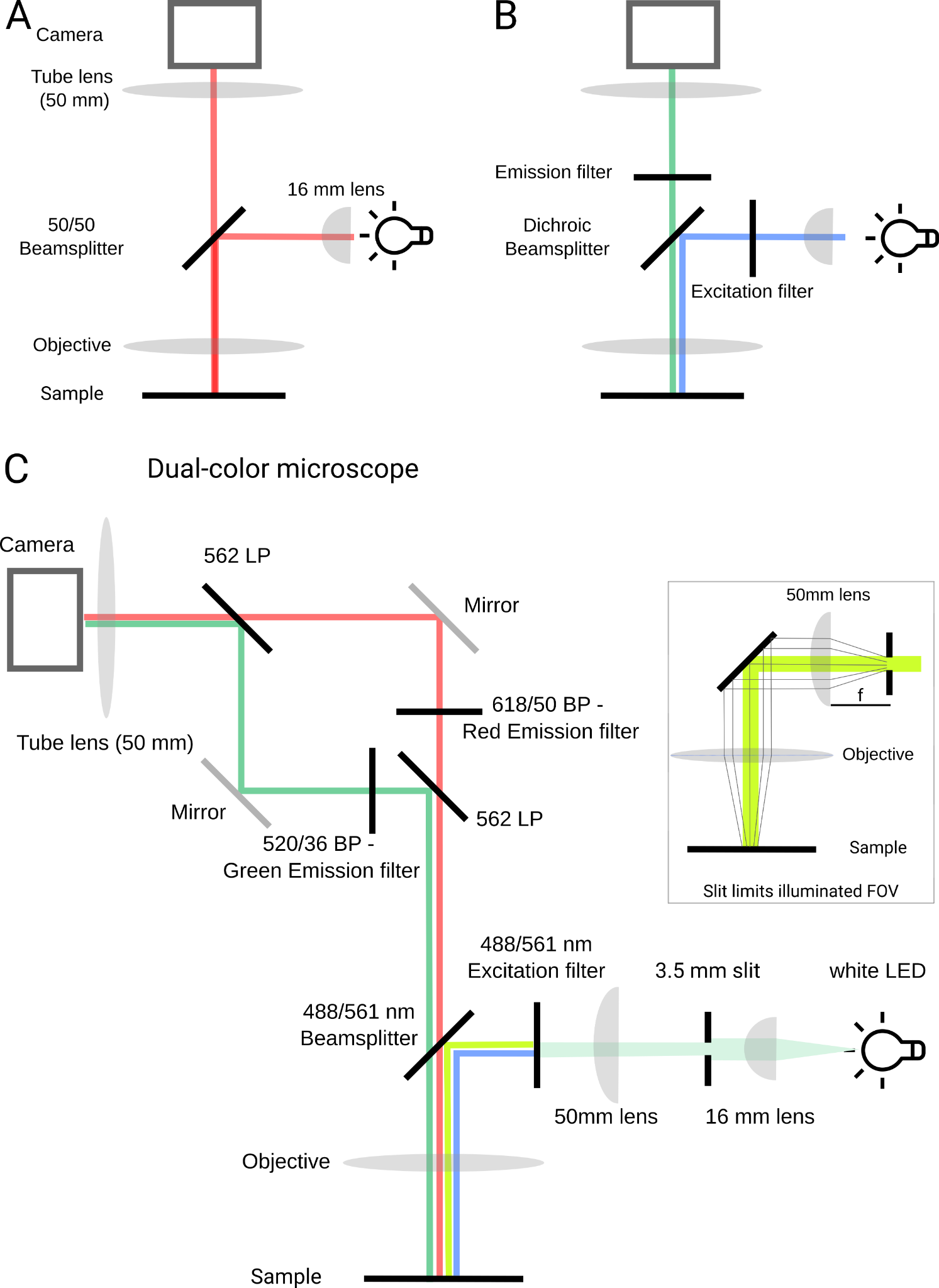

**Supplementary figure 2. Lightpath of the three different modular designs** (A) Brightfield tracking microscope. (B) Epi-fluorescence microscope. Only parts different from (A) are labeled. (C) Dual-color epi-fluorescence microscope. The inset shows the slit projection onto the field-of-view, allowing the emission to be separated onto the camera without overlap.

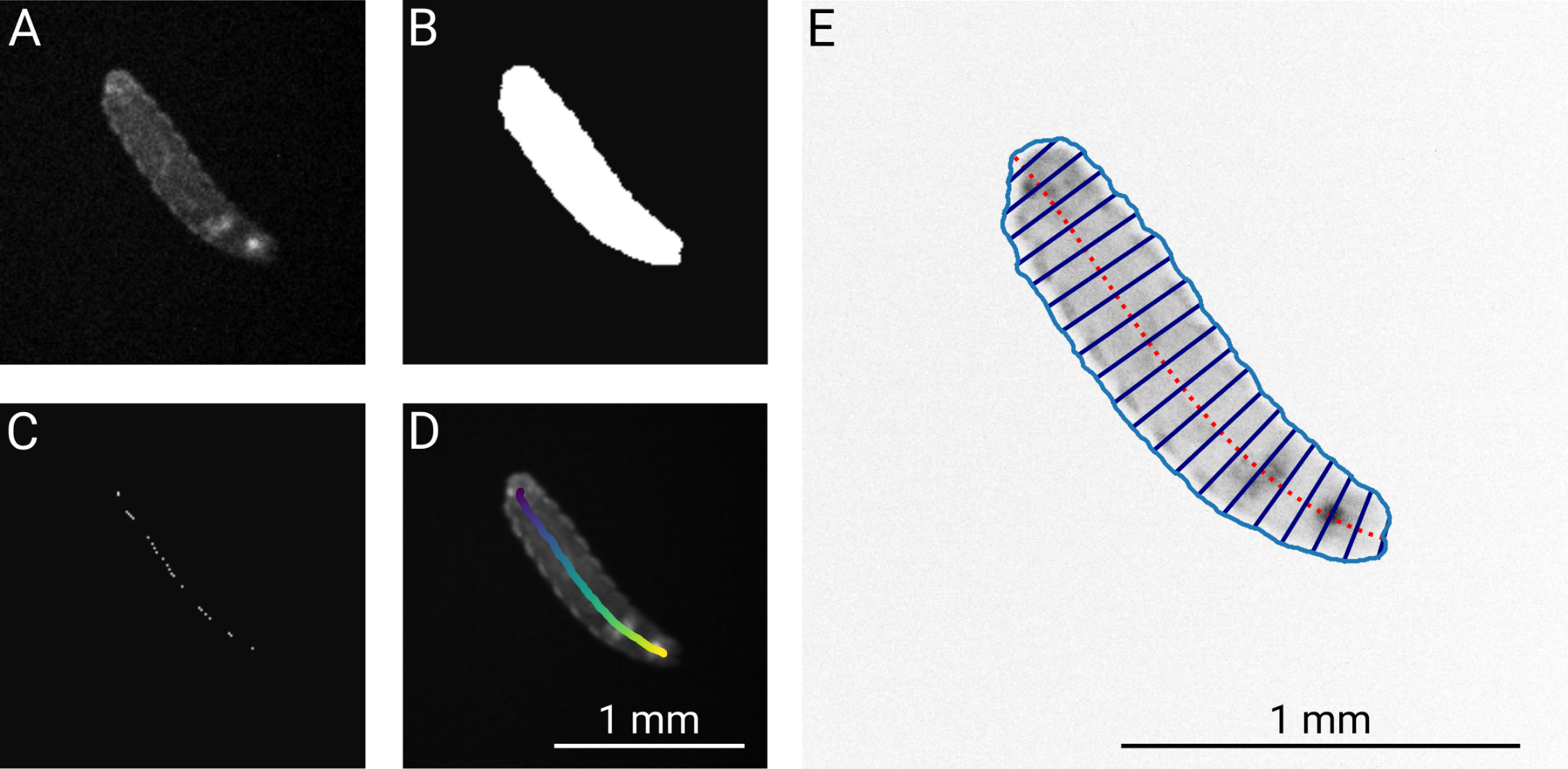

**Supplementary Figure 3. Analysis of crawling *Drosophila* larvae**

The mCherry channel of the larvae in (A) was used for segmentation (B). (C) Skeletonization and ordering of the resulting midline points was used to fit a centerline (E, red dashed line).

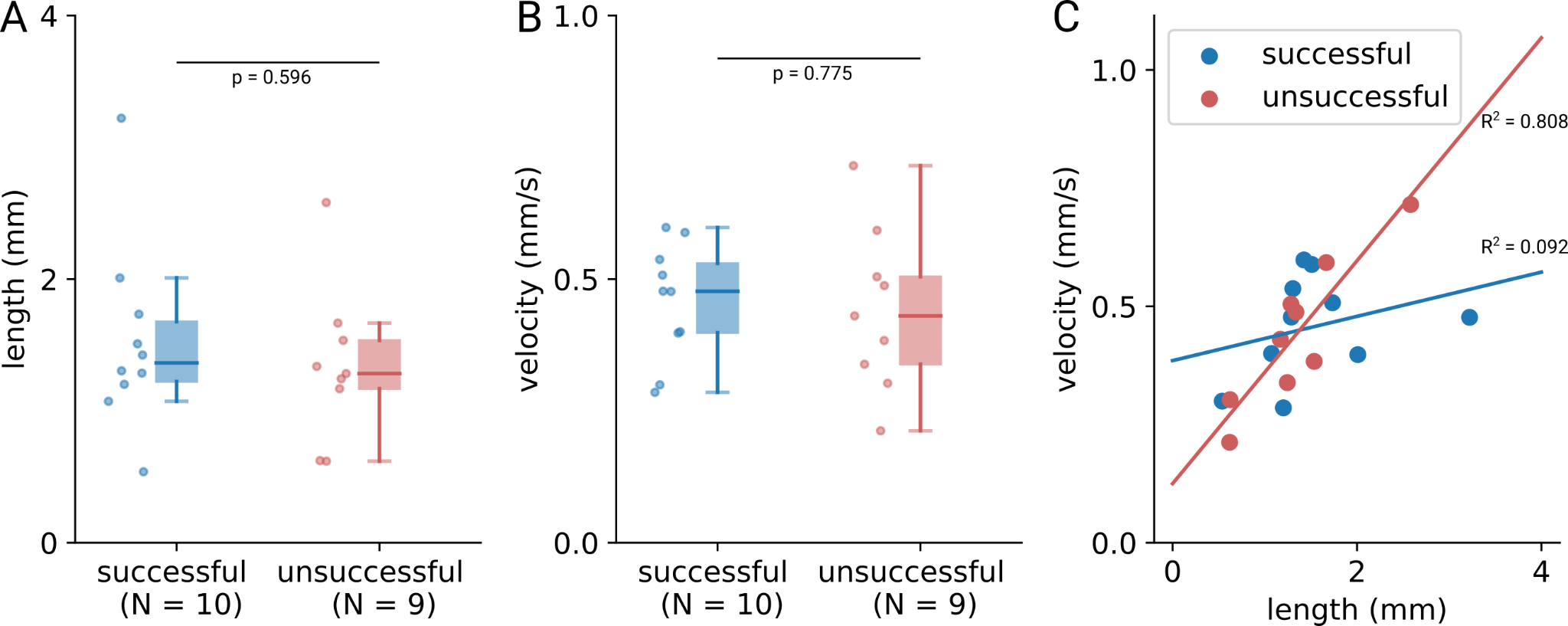

**Supplementary Figure 4. Comparison of size and velocity for successful and unsuccessful chemotaxis trials.**

(A) Length and (B) velocity of larvae that were unsuccessful (red) or successful (blue) in odortaxis trials. (C) The velocity correlated with length for unsuccessful trials, but not for successful trials. Number of animals is given in (A, B), and significance was assessed using the two-sided Mann-Whitney U-test.
